## Supplemental figures for "Aspirin-triggered resolvin D1 reduces parasitic cardiac load by decreasing inflammation through N-formyl peptide receptor 2 in a chronic murine model of Chagas disease"

Supporting Information:

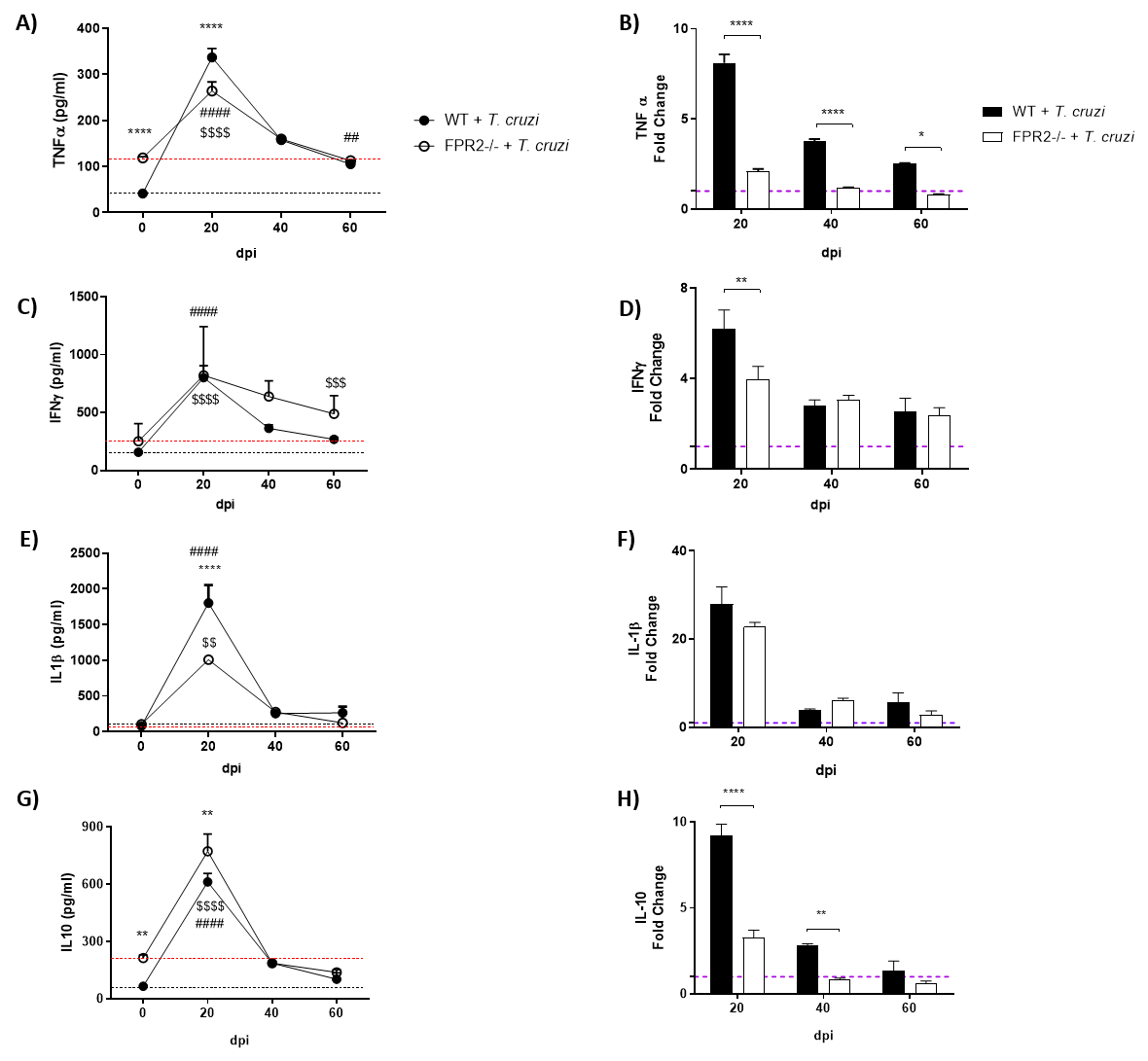

S1 Fig. Levels of cytokines in mice infected with Trypanosoma cruzi at 20, 40, and 60 dpi. The concentrations of TNFα (A-B), IFNγ (C-D), IL-1β (E-F), and IL-10 (G-H) were quantified in the serum from C57BL/6 mice uninfected and infected with T. cruzi at 20, 40, and 60 dpi, using ELISA assays. Data are expressed as the mean ± SEM from one experiment (n=5). Two-way ANOVAs and Tukey's post-hoc tests were performed to identify significant differences. Asterisks indicate significant differences between infected WT and infected FPR2. Dollar signs indicate significant differences between healthy FPR2-/- and infected FPR2-/-. Number signs indicate significant differences between healthy WT and infected WT. one symbol, p ≤ 0.05; two symbols, p ≤ 0.01; three symbols, p ≤ 0.001; four symbols, p ≤ 0.0001. dpi, days post-infection

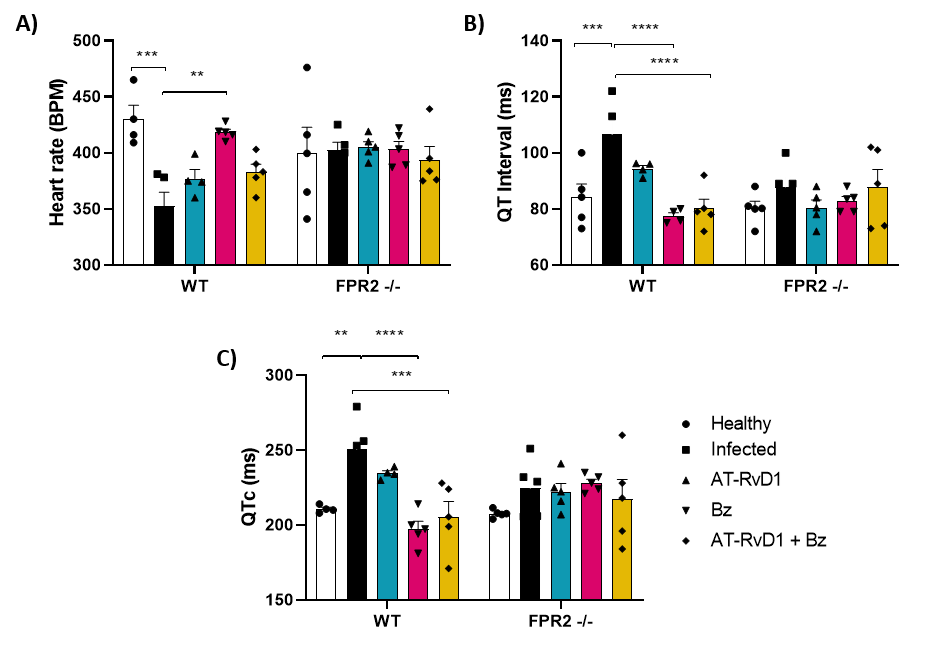

S2 Fig. Effect of AT-RvD1 and benznidazole in C57BL/6 WT and FPR2-/- mice infected with Trypanosoma cruzi on the cardiac electrical conduction system. The variation in heart rate (A), QT Interval (B), and QTc (C) are presented. The statistical analysis used was two-way ANOVA followed by Tukey's post-hoc tests (n=5). **p ≤ 0.01, ***p ≤ 0.001, ****p ≤ 0.0001. QTc, corrected QT interval.
